# Protein-level assembly increases protein sequence recovery from metagenomic samples manyfold

**DOI:** 10.1101/386110

**Authors:** Martin Steinegger, Milot Mirdita, Johannes Söding

## Abstract

The open-source *de*-*novo* Protein-level assembler Plass (https://plass.mmseqs.org) assembles six-frame-translated sequencing reads into protein sequences. It recovers 2 to 10 times more protein sequences from complex metagenomes and can assemble huge datasets. We assembled two redundancy-filtered reference protein catalogs, 2 billion sequences from 640 soil samples (SRC) and 292 million sequences from 775 marine eukaryotic metatranscriptomes (MERC), the largest free collections of protein sequences.

---

A major limitation of metagenomic studies is that often a large fraction of short reads (80% — 90% in soil [1]) cannot be assembled into contiguous sequences (contigs) long enough to allow for the prediction of gene and protein sequences. Because low-abundance genomes are difficult to assemble, the unassembled reads contain a disproportionately large part of the genetic diversity and probably an even greater share of biological novelty, which is mostly lost for subsequent analyses.

To decrease this loss and be less dependent on reference genomes, gene-centric approaches have been developed. Assemblies of hundreds of samples from one environment are pooled, genes in the contigs are predicted and clustered at ∼ 95% identity into gene catalogs [2-4]. Gene abundances in each sample are found by mapping reads to the reference gene clusters. In this way, the functional and taxonomic composition of metagenomic samples and their dependence on environmental parameters can be studied. Also, genome-based analyses are enabled by abundance binning, which finds sets of catalog genes with correlated abundances across many samples and hence likely to belong to the same genome [5, 6].

State-of-the-art assemblers for metagenomic short reads [7-9] find contigs as paths through a de-Bruijn graph. This graph has a node for each *k*-mer word in the reads and edges between *k*-mers occurring consecutively in a read. On metagenomic data, de-Bruijn assemblers suffer from a limited sensitivity-selectivity trade-off: *k*-mers have to be long and specific to avoid the graph exploding with false edges. But long *k*-mers lack sensitivity when intra-population diversities are high and overlapping reads often contain mismatches due to single nucleotide polymorphisms (SNPs). Whatever *k* is chosen, *k*-mers will be too short to be specific enough in genomic regions conserved between species and too long to be sensitive enough in regions of high intra-population diversity. This dilemma leads to short, fragmented assemblies.

Most SNPs in microbial populations lead to no change or conservative substitutions in the encoded protein sequences (**Supplementary Fig. 1**). ORFome [10] and SFA-SPA [11] therefore proposed to assemble protein instead of nucleotide sequences. But they are too slow to run on large metagenomes 40 and as de-Bruijn assemblers they suffer from the same limited specificity-sensitivity trade-off.

In addition to avoiding mismatches, assembling protein sequences also circumvents the major issue in genome assembly, sequence repeats, because proteins have much fewer and 45 shorter repeats. Furthermore, chimerical assemblies between similar protein sequences (say ≥ 97% sequence identity) are much less problematic in that they do not lead to false conclusions about which genes occur together in a genome. Therefore, protein-level assembly also increases coverage by assem-50 bling sequences that cannot be assembled on the nucleotide level due to the risk of chimeric assemblies.

Plass uses a novel graph-free, greedy iterative assembly strategy (Fig. 1) that, together with its linear-time all-versus-all overlap computation (steps 2-4) [12], scales linearly in run-55 time and memory. This permits the overlap-based assembly of huge read sets on a single server. Most importantly, by computing full alignment overlaps instead of only *k*-mer matches, Plass overcomes the specificity-sensitivity limitation of de-Bruijn assemblers, allowing it to recover several times 60 more proteins sequences from complex metagenomes.

**FIG. 1.**
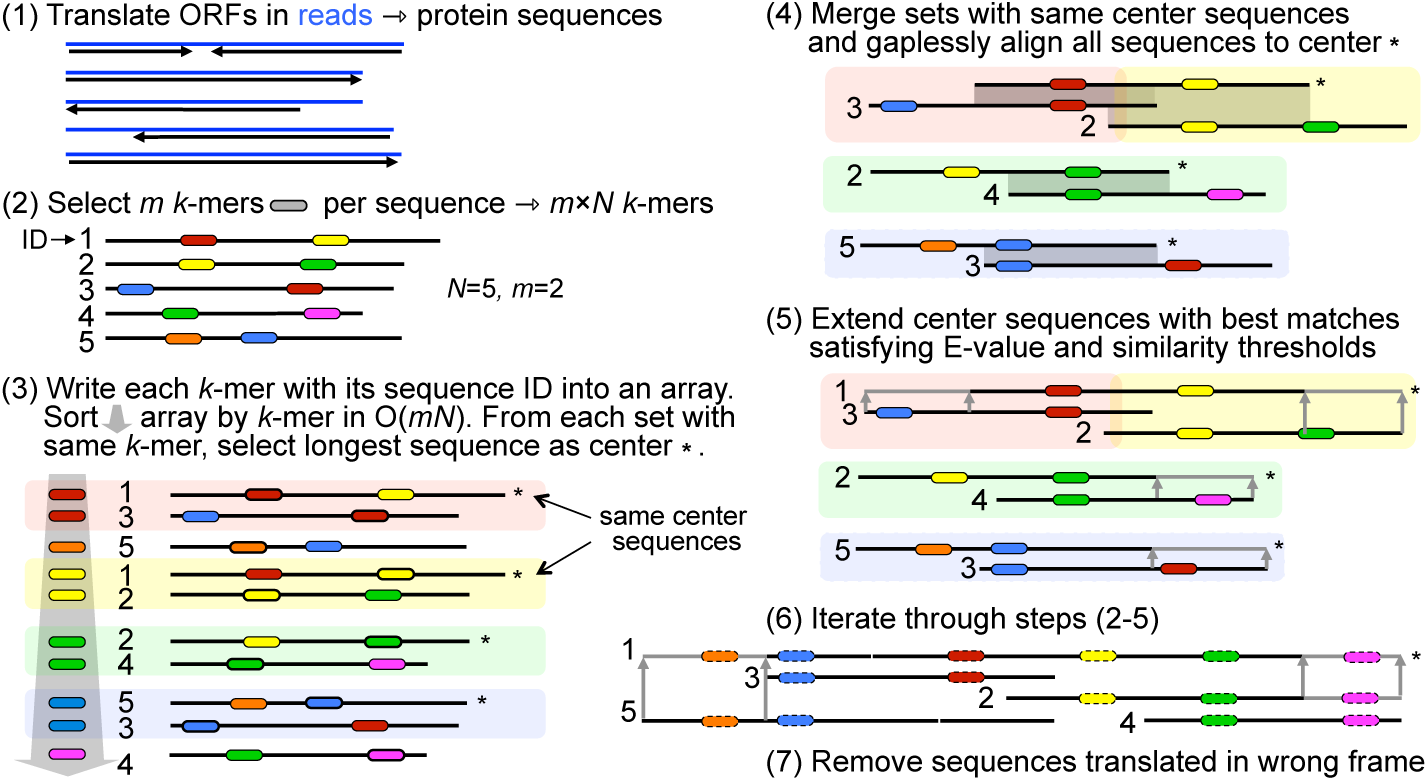
Plass workflow. (1) Merge overlapping read pairs and translate all potential ORFs with ≥45 codons into protein sequences. (2) For each of these *N* sequences, select the *m k*;-mers with the lowest hash values (default: *m =* 60, *k =* 14, reduced alphabet size = 13). Write the *mN k*-mers into an array together with sequence identifiers. (3) Sort the array by A;-mer to find for each *k*-mer the set of sequences containing it, and assign the longest sequence as the set’s center. (4) Resort the array by center sequence into groups and gaplessly align the center sequence to each group member (< *mN* alignments). Remove sequences with insufficient *E*-value (default: > 10^−5^) from the group. (5) Iteratively extend each center sequence by the remaining group member with highest sequence identity (default: ≥ 90%) until all group members have been processed. (6) Iterate steps 2 to 5 (default: 12 times). (7) Remove sequences translated in the wrong frame using a neural network.

Plass needs to keep the protein sequences in main memory to avoid random disk access (step 4). It therefore needs 1 byte of memory for every amino acid translated from the input reads, or ∼ 500 GB RAM to assemble 2-3 billion 2 x 150 bp reads. In comparison, memory requirements and runtimes of overlap graph assemblers scale superlinearly with the number of reads. Plass therefore combines the high specificity-sensitivity of overlap graph assemblers with the linear runtime and memory scaling of de-Bruijn graph assemblers.

Due to our greedy assembly approach, the most critical aspect to analyze is what fraction of the sequences are wrongly assembled (precision). To challenge Plass, we sought two hard datasets containing many related genomes, as this increases the risk of chimeric assemblies [13].

The first set consists of 96 single-cell assembled genomes of *Prochlorococcus* [14] taken from a single sample of seawater. These cyanobacteria are known for their high intra-species genetic diversity. The second, very hard set contains 738 singlecell assembled genomes, 489 of which are *Prochlorococcus*, are *Synechococcus*, and 199 are genomes from a diverse range of prokaryotic and viral groups [15].

As ground truth reference we predicted protein sequences on the genomes using Prodigal [16]. We simulated 2 x 150 bp reads with a mean coverage of 1 for each genome.

We assembled protein sequences from these nucleotide reads using Plass and SFA-SPA [11]. We assembled nucleotide contigs with three widely used nucleotide assemblers, Megahit [8], metaSPAdes [9], and Velvet [7], the first two of which were among the top assemblers in recent benchmarks [13, 17, 18]. We predicted protein sequences in their contigs using Prodigal and, to ignore unassembled reads, we removed protein sequences with less than 100 residues.

The assembly sensitivity is the fraction of amino acids in the reference proteins that have a sequence match with at least *X%* sequence identity with an assembled sequence. To avoid giving too much weight to highly conserved proteins, we redundancy-filtered the reference proteins for the sensitivity analysis using Linclust [12] with 95% sequence identity. The sensitivity is similar for the three nucleotide assemblers, whereas Plass assembles up to 56% more residues correctly than the next best tool, metaSPAdes **(Fig. 2a, top)**.

**FIG. 2.**
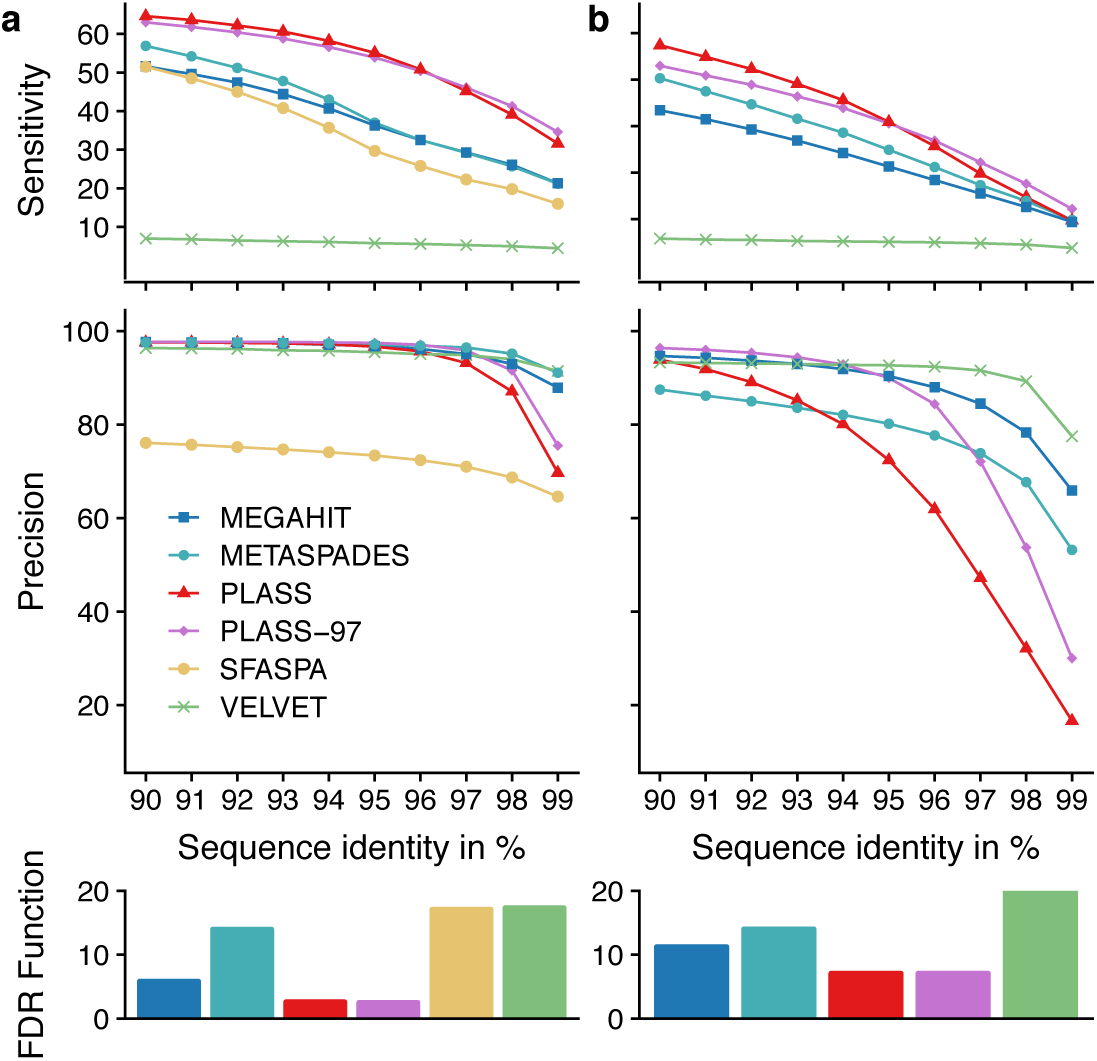
Plass assembles many times more protein sequences from various environments than the state of the art. **(a)** Sensitivity and precision of protein sequences assembled from synthetic reads sampled **(a)** from 96 assembled genomes of single *Prochlorococcus* cells [14] and **(b)** from 738 single-cell assembled genomes of diverse marine prokaryotes and viruses [15]. For the three nucleotide assemblers we predicted protein sequences on their assembled contigs with Prodigal [16]. Top: Sensitivity is the fraction of reference sequence amino acids that have are aligned to an assembled protein sequence with a sequence identity at least the value on the x-axis. Middle: Precision is the fraction of assembled amino acids that are aligned to a reference protein with sequence identity at least the value on the x-axis. Bottom: False discovery rate (1 — precision) of orthology-based functional annotations of assembled proteins. Colors and order of tools as in previous legend.

The Plass-assembled proteins cover over 80% of the Megahit and metaSPAdes assemblies at 99% minimum sequence identity, whereas the latter cover only around 40% of the Plass assembly at this cut-off **(Supplementary Fig. 3)**.

The precision is the fraction of assembled amino acids that have at least *X%* sequence identity with an assembled sequence.

Since ORFs are predicted on the nucleotide assemblies using the same tool used to define the reference protein sequences on the single-cell genomes, whereas Plass uses a very different approach (Online Methods), the benchmark is biased against Plass. Nevertheless, Plass achieves the same precision below *X%* = 97%, where all assemblers except SFA-SPA achieve similar precision **(Fig. 2a, middle).** The 2% — 7% of missing precision at *X% =* 90% are mainly caused by mispredicting open reading frames (ORFs) in the assembled sequences or on the single-cell genomes.

Plass’ neural network filter (Online Methods) for suppressing proteins translated in wrong frames raises the precision at *X% =* 90% on the very hard set by a few percent **(Supplementary Fig. 3b).**

Plass filters out sequences translated from wrong ORFs based on their amino acid and dipeptide composition, which differs from correctly translated, real protein sequences. We-trained a neural network using as features the201ength-normalized amino acid frequencies of the sequence and the621ength-normalized dipeptide frequencies in a reduced alpha-bet of size6. Our fully connected network has56input nodes,a hidden layer of96nodes, and a single output node.

However, Plass produces much fewer proteins at 99% sequence identity than the nucleotide assemblers, particularly on the very hard data set **(Fig. 2b).** Increasing the sequence identity threshold for merging sequence fragments from 90% to 97% (pink trace) markedly improves sensitivity and precision on both datasets, but still much fewer proteins match a reference protein with identity of ≥ 97%.

To test the impact of the assembly of chimeric protein sequences on the quality of functional annotations, we annotated each protein sequence assembled from the simulated reads and each reference protein from the single-cell genomes with an orthologous group using the eggNOG mapper [19]. We compared the eggNOG annotation of each assembled protein sequence with the annotation of the best-matching sequence found in the reference protein set and scored a true positive (TP) if annotations matched and a false positive (FP) otherwise. Assemblies that cannot be assigned to a reference are FPs. Despite Plass’ lower assembly precision, annotations of its proteins achieve lower false discovery rates (FP/(TP + FP)) than those of the other assemblers, on both data sets **(Fig. 2, bottom).** We believe this is due to (1) the high conservation of molecular and cellular functions at sequence identities > 90% (above 60% identity, 90% of proteins conserve of all four EC number digits [20]), (2) the limited ability of homology-based function annotation tools to predict the effects of point mutations, and (3) the positive impact of more complete protein sequences on the prediction accuracy.

On real metagenomic datasets, no ground-truth set of reference sequences exists. Therefore precision cannot be measured, but sensitivity in terms of the total number of assembled amino acids can be compared. We used four representative test sets: a single 11.3 Gbp sample from the human gut [21], 775 samples with 15Tbp of eukaryotic metatranscriptome reads from TARA [22], a 31 Gbp sample from Hopland grass soil *(Brodie et al.,* unpublished), and 538 Gbp of reads in 12 samples from the same project to test the benefits of co-assembly (Fig. 2b-e). All datasets contain 2 x 150 bp overlapping paired-end sequences, except the metatranscriptomics sample, which has 2 x 102 bp reads.

We compared the Marine Eukaryotic Reference Catalog (MERC) assembled by Plass to the Marine Atlas of Tara Oceans Unigenes (MATOU) [22] assembled by Velvet. The gut and soil datasets could not be assembled with Velvet due to insufficient memory, and we could only compare Plass to Megahit and metaSPAdes. The twelve Hopland soil datasets with 1.5 billion reads could be co-assembled in one go by Plass. Megahit raised an out-of-memory exception, therefore we assembled each sample separately and pooled the contigs. For human gut, Hopland soil, and the soil co-assembly, Plass took 4h 20min, 6h 20min and 360h respectively, while Megahit took 3h, 21h 30min and 200h respectively.

On the gut sample, Plass assembled 32% more amino acids than Megahit/Prodigal at a length cut-off of 100 amino acids; (**Fig. a-d**, top). The marine eukaryotic reference catalog (MERC) assembled by Plass is 2.8-fold larger than MATOU assembled by Velvet, and on the Hopland soil data Plass assembled 2.7 times more than Megahit. In the soil co-assembly, Plass co-assembled 10 times more amino acids than the pooled assembly of Megahit. The increase of the ratio with sequence length in the top of Fig. 2d,e indicates that the sequences assembled by Plass are significantly longer than those of Megahit/Prodigal. These gains in recovered protein sequences are similar at all levels of redundancy up to 80% se; quence identity (bottom half of Fig. 2b-e).

We wanted to know how strongly the improved sensitivity of Plass affects the apparent taxonomic composition. We implemented the 2bLCA protocol (**Supplementary Fig. 4**) [23] to map each read via its translated ORF to an assembled protein sequence and each protein sequence to a node in the taxonomic tree. By transitivity this maps reads to taxonomic nodes. The absolute number of reads mapped to various taxonomic nodes (**Supplementary Fig. 5a**) is around twice higher for Plass than for Megahit/Prodigal. Remarkably, the distribution over taxa can deviate quite substantially between the assembly methods (**Supplementary Fig. 5b**), which could be caused be a systematic dependence of the assembly sensitivity on genomic coverage (**Supplementary Fig. 5c**).

Plass is well suited to large-scale applications. We assembled a Soil Reference protein Catalog (SRC) from 18 Tbp of reads from all 640 soil samples that were sequenced between 01/2016 and 02/2018 using Illumina HiSeq or NovaSeq with 2 × 150 bp paired-end reads. Each sample was assembled on a server with 2 × 8 cores and 128 GB memory, resulting in 12 billion protein sequences after a total runtime of about six weeks on 25 servers. We clustered the sequences to 90% sequence identity at 90% minimum coverage using Linclust [12], resulting in 2 billion sequences with an average length of 163 amino acid residues. Among those, at least 52.3 million sequences are complete, meaning that Plass found the stop codon and the earliest possible start codon (Online Methods). This dataset contains 6.8, 4.0 and 3.9 times more amino acids than the Uniprot database after redundancy-filtering both databases at 90%, 70% and 50%, respectively (Fig. 3a).

**FIG. 3.**
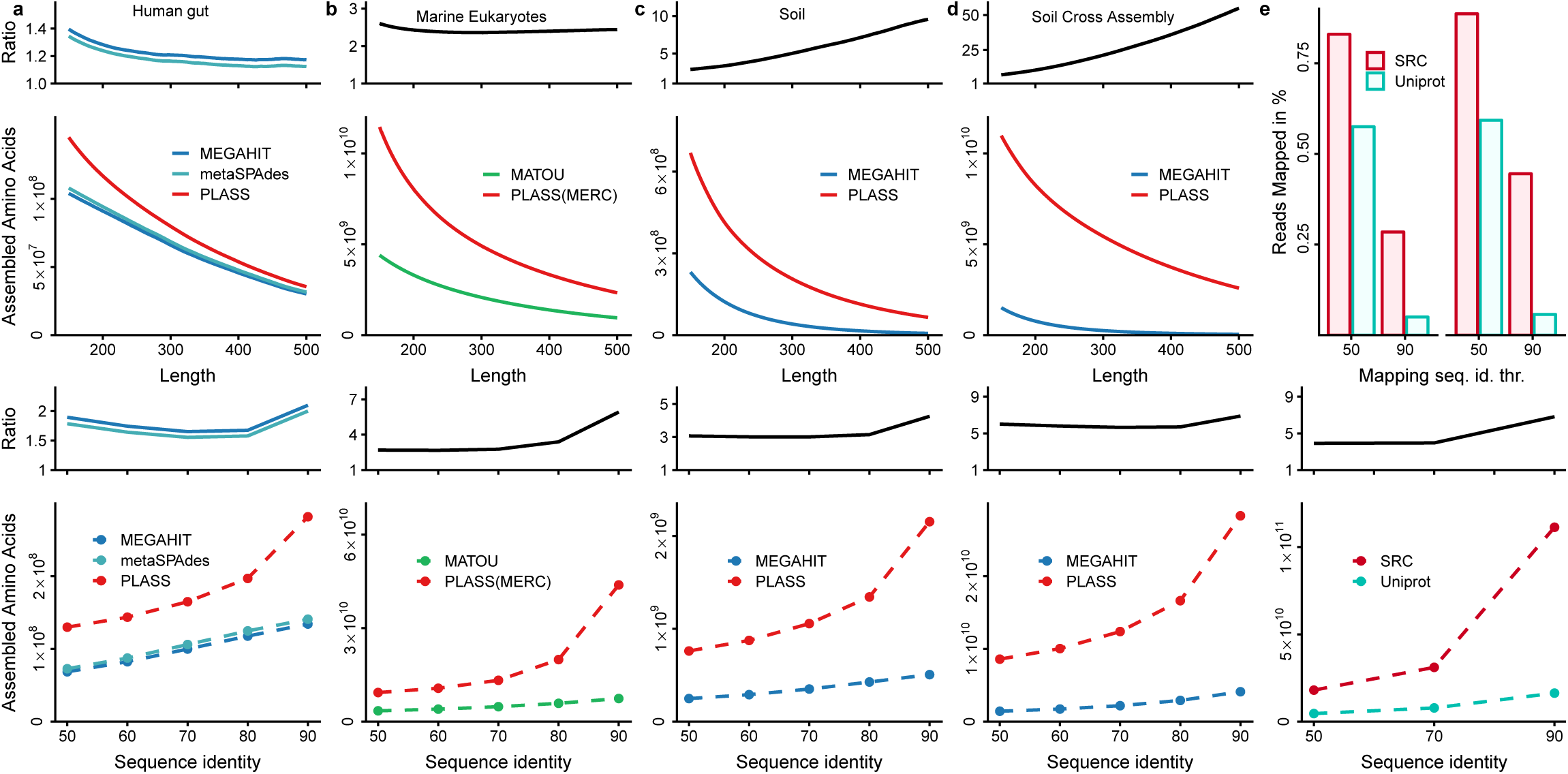
Plass assembles many times more protein sequences from various environments than the state of the art. (**a-d**) Total number of amino acids in redundancy-filtered sets of protein sequences assembled by Plass (red traces) compared to the total number of amino acids of redundancy-filtered protein sequences predicted by Prodigal on contigs assembled by Megahit (**a,c,d**: blue) or on contigs in the eukaryotic metatranscriptomes reference assembly (b: green) [22]. Top half: dependence on the minimum protein sequence length using a redundancy-filtering with 80% maximum pairwise sequence identity. Bottom half: dependence on the strength of redundancy filtering for a minimum sequence length of 100 amino acids. Black traces: fold increase in total assembly length by Plass versus the state of the art. (**e**) Top half: Fraction of reads sampled from two soil metagenomes (not included in the SRC) that could be mapped with 90% and 50% minimum sequence identity to a sequence in the SRC or in Uniprot. Bottom Half: Numbers of amino acids in SRC (red) and Uniprot (turquoise) and their ratio (black) after redundancy-filtering with a maximum pairwise sequence identity of 50%, 70% and 90%.

To assess the degree to which the SRC represents the diversity of soil metagenomes, we selected two soil samples not used for building the SRC, randomly sampled and merged 10 000 overlapping 2 × 150 bp read pairs, predicted protein sequences with Prodigal, and searched with these through the 90%-redundancy-filtered versions of the SRC and the Uniprot, using the map workflow of MMseqs2 [24]. Fig. 3b,c shows the fraction of reads that obtained matches with at least 50% and 90% sequence identity. At 50% threshold, 82.5% and 89.5% of the soil reads matched to the SRC in the two samples, while only 62% and 64% matched to the Uniprot.

The chief limitation of Plass is that, unlike nucleotide assemblers, it cannot place the assembled protein sequences into genomic context. Furthermore, Plass relies on six-frame translation. It therefore cannot assemble intron-containing eukaryotic proteins from metagenome data, although, as shown, it can assemble eukaryotic proteins from transcriptome data. Another important drawback is its inability to resolve homologous proteins with sequence identities above ∼ 95%, for example originating from closely related strains or species. However, in our tests this had little impact on the accuracy of predicted functions (Fig. 2). Also, one can argue that bacterial phenotypes are determined mainly by the complement of horizontally acquired accessory genes such as virulence factors, much more so than by minor variations in protein sequences. Whereas Plass is clearly worse than nucleotide assemblers in resolving such variation, it excels at assembling more complete protein complements of metagenomes.

In conclusion, Plass is well-suited for very large-scale metagenomic applications, for example to generate reference protein sequence catalogs for every major type of environment. In addition to facilitating metagenomics analyses, these catalogs can be mined for proteins of interest to biotechnology or pharmacology. They will also improve homology detection, protein function annotation and protein structure prediction [25] by enriching multiple sequence alignments with diverse homologs.

## Methods

Methods, including statements of data availability and references, are available in the online version of the paper.

## Supporting information

## Acknowledgements

We are grateful to Cedric Notredame and Chaok Seok for hosting MS at the CRG and SNU for 12 and 30 months, respectively. We thank Shinichi Sunagawa, Folker Meyer and Alex Sczyrba for helpful discussions and Titus Brown for his early analysis and detailed feedback on Plass results. This work was supported by the EU’s Horizon 2020 Framework Programme (Virus-X, grant 685778).

## Author contributions

MS & JS designed research, MS & MM developed code and performed analyses, MS, MM & JS wrote the manuscript.

## ONLINE METHODS

Plass proceeds in seven steps summarized in Fig. 1.

### Merging paired-end reads and ORF calling

Longer reads increase the precision and sensitivity of the assembly due to longer overlaps obtaining higher statistical significance. In step 1, Plass therefore merges overlapping paired-end reads into longer sequences using code from the open-source FLASH tool [23], which we integrated into Plass (step 1**a**).

Furthermore, in step 1, Plass extracts all open reading frames (ORFs) with at least 45 codons and translates them into protein sequences. (Alternative codon tables can be specified with option `-translation-table`.) To determine the correct start codon later on, it also extract and translates all ORFs with at least 20 codons starting with a putative `ATG` start codon that is the first `ATG` codon after a stop codon in the same frame. Because the coding sequence cannot start before such an `ATG`, these sequences help Plass to predict start codons later on (see “Predicting start codons” and **Supplementary Fig. 6**).

### Finding overlaps in linear time

The identification of all overlapping alignments (Fig. 1, steps 2-4) is critical for the performance of overlap assemblers. Previously proposed protein-level assemblers have a runtime complexity that scales quadratically with the input set size [24,25]. A typical metagenomic read set with 100 million reads requires 10^16^ comparisons with a quadratic method. To speed up the computation, we adapted our linear-time clustering algorithm Linclust [26] for assembly.

In step 2, Plass transforms each protein sequence into a reduced amino acid alphabet, whose 13 letters represent the following groups of amino acids: (L, M), (I, V), (K, R), (E, Q), (A, S, T), (N, D), and (F, Y). From each reduced sequence it selects m (default: *m* = 60) *k*-mers (or *l — k +1* if the sequence of length *l* contains only *l − k + 1 < m k*-mers). The selected *k*-mers are those with lowest hash values. Our rolling hash function [26] maps each *k*-mer onto a range of [0,2^16^] such that even single residue changes result in quasirandom, unrelated hash values. For each of the ∼ *mN* selected *k*-mers, Plass stores in an array the *k*-mer index (8 bytes), the sequence identifier (4 bytes), the *k*-mer position in the sequence (2 bytes) and the length of the sequence (2 bytes).

In step 3, Plass sorts the array by *k*-mer index and sequence length to find the sets of sequences containing the same *k*-mer. For each set, it picks the longest sequence as the center sequence. For each member of the *k*-mer set, it overwrites the *k*-mer index with the center sequence identifier and computes the diagonal *i — j* on which the shared *k*-mer match occurs, where *i* is the *k*-mer position in the center sequence and *j* is the position in the member sequence. The array now contains the center sequence identifier, the member sequence identifier, the *k*-mer match diagonal, and the length of the member sequence. It sorts the array again, this time by center sequence identifiers, and removes duplicate center-member pairs. If more than one diagonal match between a center and member sequence is found only the match with lowest diagonal is kept.

In step 4, Plass computes an ungapped local alignment between each center sequence and each group member, using onedimensional dynamic programming on the diagonal *i − j* of the *k*-mer match. It computes E-values using ALP [27] and, by default, the Blosum62 substitution matrix. Alignments with an E-value > 10^−5^ (default) and a sequence identity < 90% (default) are rejected.

### Extending protein reads

In step 5, Plass extends the center sequence by concatenating the non-overlapping residues of the member sequence with highest similarity in the overlap. More precisely, it processes the list of alignments with the member sequences in order of descending overlap sequence identity, until one side of the center sequence has been extended and the other side has either been extended as well or has no extending alignments left in the list. Then it realigns the extended center sequence with all yet unprocessed member sequences and iterates the extension until the entire list of alignments has been processed.

### Iterative assembly

Plass iterates through steps 2 to 5 twelve times (default), each time updating the original version of the center sequences with their extended versions and keeping all other sequences unchanged (step 6). To extract different *k*-mers in each new iteration, we increment the step size of the circular shift inside our rolling hash function [26].

### Removing proteins translated in wrong frames

In step 7, Plass removes sequences translated from wrong ORFs or assembled from such sequences. ORFs translated in the wrong frame contain a stop-codon approximately every 64/3 ≈ 21 residues, and so only a fraction of around exp(—45/21.3) ≈ 12% contain ≥ 45 codons.

Plass filters out sequences translated from wrong ORFs based on their amino acid and dipeptide composition, which differs from correctly translated, real protein sequences. We trained a neural network using as features the 20 length-normalized amino acid frequencies of the sequence and the 6^2^ length-normalized dipeptide frequencies in a reduced alphabet of size 6. Our fully connected network has 56 input nodes, a hidden layer of 96 nodes, and a single output node. We trained the network using the Keras deep-learning framework using the Adam optimizer with a 10% drop-out probability and the binary cross-entropy loss function. We leave 10% of the data out for cross validation. The network is integrated into Plass using Kerasify.

To train the network, we created a positive set of known coding sequences and a negative set of sequences translated in a wrong reading frame. The positive set contained 2.4 million proteins sampled from the prokaryotic subset of the Uni-clust30 representative sequences [28]. For the negative set, we extract all ORFs from 757 prokaryotic genomes contained in the KEGG database [29] and clustered them using MMseqs2 [30] with a maximum sequence identity of 30% and a minimum coverage of 80%. Clusters without any member with coding sequence annotation in KEGG or homology to entries in Uniclust30 (requiring an *E*-value of < 10^−3^) were extracted. From these, we sampled 2.4 million sequences.

### Predicting start codons

To determine the correct start codon and minimize overextension at the N-terminus in the ORF translation step 1, Plass marks with a prepended asterisk * those methionine residues that represent the first `ATG` after a stop codon in the same frame, as this implies that the coding sequence can at the earliest start there (**Supplementary Fig. 6**).

After the alignment and extension step 4 in the first iteration, Plass reconstructs the multiple sequence alignment of all merged sequences. Where at least 20% of all methionines in a column are marked by a prepended asterisk, it removes the preceding residues from all other sequences and prepends an asterisk to all sequences to mark the start. If several columns fulfill the 20% criterion, it trims the sequences at the most downstream of these columns. The start codon prediction is only done in the first iteration to save time and disk space.

### Suppressing repetitive sequences

Protein repeats can lead to unwanted extensions during assembly. We therefore detect sequences with repeat regions during step 2 as those containing at least 8 (default) identical *k*-mers (in the 13-letter alphabet). These sequences are ignored during all steps.

### Memory-efficient processing of huge input sets

Plass needs 1 byte per residue of translated protein sequence generated in step 1 to keep these sequences in memory and avoid random disk accesses in the alignment step 4. But as we saw, the *k*-mer array in steps 2 and 3 occupies *m* × 16 bytes = 720 bytes of memory per sequence, which is around 16 times more than 1 byte per residue. We removed this bottleneck, for systems with insufficient main memory, by splitting the *k*-mer array into a number of chunks and processing them sequentially, with little loss in speed.

We compute the maximum number of chunks S that still allows one chunk to fit into the available system memory *M* as *S* = ⌈*mN x* 16 byte/*M*⌉. For each chunk c out of S, we proceed with steps 2 and 3 exactly as described before except that we only extract *k*-mers whose index *R* satisfies (*R* mod *S*) = *c* and that we store the chunks of *k*-mer arrays on hard disk. After all splits have been computed, we merge them into a single *k*-mer array.

### Assembly quality benchmark

We could not use the standard benchmark developed by the Critical Assessment of Metagenomic Interpretation (CAMI) [31], because the mutation model of the sgEvolver tool used for simulating population microdiversity (strain diversity) does not penalize frame-disrupting indels and non-conservative substitutions within coding regions. This leads to very low and unrealistic conservation of coding regions. The synthetic reads generated with such a model are certainly realistic enough to test nucleotide assemblers but would render protein-level assembly absurdly unsuitable.

We sought to construct a genomic benchmarking set that would contain a high degree of *natural* variation, in which the genomic sequences reflect the actual evolutionary pressures on them.

We downloaded from the sequence read archive (SRA) at the NCBI/NIH two sets of genomes assembled from single cell sequencing libraries. The first set contains genomes of 96 *Prochlorococcus* genomes [32].

These cells were taken from the same ocean water sample and represent a population of the cyanobacteria *Prochlorococcus*, the most abundant marine photosynthetic organism on earth noted for high intra-species diversities. Sequence identities of 16S rRNA ITS sequences in a matched sample are between 50% and 100%.

The second set contains 738 single-cell genome assemblies (NCBI project PRJNA445865) consisting of 489 *Prochlorococcus,* 50 *Synechococcus,* 82 SAR11, 17 SAR116, 16 SAR86, 9 extracellular virus particles, and 75 additional sympatric microorganisms, sampled at 22 locations in the Atlantic and Pacific Oceans [33].

As ground truth reference we predicted protein sequences on the genomes using Prodigal [34] and removed sequences shorter than 100 residues, resulting in a redundant reference set of 109 014 protein sequences for set 1 and 829 899 for set 2. We reduced the redundancy by clustering with Linclust at 95% sequence identity and 99% minimum coverage of the shorter sequence (options `--cov-mode 1 −c 0.99 --min-seq-id 0.95`), resulting in the non-redundant reference set with 14 943 (set 1) and 460 653 (set 2) sequences.

We created two synthetic read data sets from the two sets of single-cell genomes, setting the mean coverage to 1 for each genome, which yielded 392 790 reads for set 1 and 4 994 546 for set 2. We used `randomreads.sh` from the BBmap software suite with options `paired snprate=0.005 adderrors coverage=1 len=150 mininsert=150 maxinsert=350 gaussian=true` to simulate 2 × 150 bp paired-end overlapping reads with sequencing errors.

We then assembled the synthetic paired-end read data set with Megahit, metaSPAdes, Plass, SFA-SPA and Velvet, using the default parameters of each tool. We also tested Plass with with a stricter minimum sequeunce identity for merging sequences (option `--min-seq-id 97`, “Plass-97” in Fig. 2. For the nucleotide assemblers, we called proteins from the assembled contigs using Prodigal in metagenomics mode. We ignored all proteins shorter than 100 residues.

We calculated the *precision* by searching with the assembled proteins through the redundant reference set, using MMseqs2 with options `-a −s 5 --max-seqs 5000 --min-seq-id 0.89`. We filtered the aligned set by minimum sequence identity thresholds between 90% and 99%. For each search result, we only considered the longest alignment that fulfills the minimum sequence identity criterion. We computed the precision for each sequence identity threshold as the ratio of the total count of aligned residues divided by the total length of the assembled proteins. 100% precision is reached when all assembled protein residues can be aligned to a reference protein sequence.

We calculated the *sensitivity* by searching with the nonredundant reference set through the assembled proteins, using MMseqs2 with options `-a −s 5 --max-seqs 500000 --min-seq-id 0.89`. We filtered the aligned set by minimum sequence identity thresholds between 90% and 99%. For each search result, we only considered the longest alignment that fulfills the minimum sequence identity criterion. We computed the sensitivity for each sequence identity threshold as the ratio of the total count of aligned residues divided by the total length of the proteins in the non-redundant set. 100% sensitivity is reached when all reference protein residues can be aligned to an assembled protein sequence.

### Accuracy of functional annotation of assembled proteins

To test the impact of chimeric assemblies and assembly quality on functional annotation quality, we measured the accuracy of functional annotations on the proteins assembled from the reads simulated from the single-cell marine genomes (Fig. 2). We functionally annotated each protein sequence assembled from the simulated reads and each reference protein from the single-cell genomes with an Orthologous Group using the eggNOG mapper[35] and the eggNOG database (version 4.5.1)[36] (options `-d bact −m diamond-override-cpu 16`). We compared the eggNOG annotation of each assembled protein sequence with the annotation of the best-matching sequence found in the reference protein set using MMseqs2. If the Orthologous Groups differ the annotation is false positive (FP), otherwise true positive (TP).

### Protein sequence recovery on metagenomic datasets

For the benchmark test on real metagenomic data (**Fig. a-d**) we used the following datasets: (b) a single human gut sample from SRA (`SRR5024285`) [37], (c) 775 samples from Tara eukaryotic metatranscriptomes downloaded from the ENA (PRJEB6609) [38], (d) a soil sample from the IMG project 1003784 (sample: 6398.7.44014), (e) 12 samples from the same project (samples: 6679.7.51457 6478.6.45123, 6679.6.51456, 6398.7.44014, 6478.7.45124, 6674.6.51288, 6679.5.51455, 6674.4.51285, 6478.5.45122, 6478.4.45121, 6674.3.51284, 6674.5.51286). The soil data is also available at the NCBI Project PRJNA330082. All samples used in Fig. 2b,d,e consist of paired-end reads of 2 × 150 bp length, while c consists of reads with 2 × 102 bp length.

We assembled paired-end reads in datasets **b,d,e** using Megahit and Plass with default parameters. The benchmarks for sets in Fig. 2b-d were carried out on a single. The coassembly in Fig. 2e was performed on a server with two 14-core Intel Xeon E5-2680 v4 CPUs with 768 GB RAM. During the co-assembly, Megahit aborted with a segmentation fault on the 768 GB server. We therefore performed twelve separate assemblies and pooled the results.

We could compare Plass only to Megahit on datasets b,d,e, since Velvet terminated with segmentation faults, metaSPAdes terminated with messages specifying a required amount of RAM in excess of the available 128 GB, and SFA-SPA did not finish execution within three days.

For Fig. 2c, we assembled the 775 Tara metatranscriptomes using Plass and compared the results with the Marine Atlas of Tara Oceans Unigene (MATOU) catalog [38], assembled using Velvet. For that purpose, we called protein sequences using Prodigal in metagenomics mode on all MATOU contigs, since these often do not contain full-length protein sequences. Eukaryotic protein sequences contain repeats more frequently than viral or prokayotic ones. We therefore masked low complexity regions of the assemblies created by Plass using tantan [39] and removed all assembled proteins with more than 50% masked residues.

To analyze the diversity of the obtained sets at various redundancy levels, we clustered all assembled protein sequence sets with Linclust using the parameters `--kmer-per-seq 80 --cluster-mode 2 --cov-mode 1 −c 0.9` at sequence identity thresholds `--min-seq-id` from 50% to 90%.

### Taxonomic classification and quantification

We investigated the influence of the assembly method on the taxonomic composition (**Supplemental Fig. 5**). Instead of matching nucleotide reads to reference genomes, we here perform the taxonomic matching on the protein level because, first, many species sampled with metagenomics do not contain a close homolog in the reference databases, and second, protein-level comparison afford a much higher sensitivity to match to more distantly related sequences.

Our strategy is to (1) map reads - via the translated ORFs they contain - to assembled protein sequences and to (2) map the assembled protein sequences to taxonomic nodes in the NCBI taxonomic tree. We thereby map transitively each read to one taxonomic node.

To map the assembled protein sequences to taxonomic nodes (step 2 above), we implemented the 2bLCA protocol [40] as new MMseqs2 module `mmseqs taxonomy` (**Supplementary Fig. 4**) and assigned the assembled protein sequences to the 90% redundancy-filtered Uniprot database (Uniclust90 2017_07) [28], which contains taxonomic assignments to the NCBI tree for each sequence.

Using the two-step transitive mapping, we computed read counts for all taxonomic nodes. We then pooled the counts for each phylum in the tree and in addition recorded counts of reads assigned by 2bLCA to taxa above the phylum level. Only the 8 most abundant taxa were then kept, and counts of all others were pooled into a category “Others".

In **Supplementary Fig. 5a** we show the results for the soil sample assemblies from c (blue: Megahit, red: Plass) and the assemblies of the 12 soil samples from **d** (light blue: Megahit, light red: Plass), together with the ratios on top. The inset gives the fraction of reads in the single and the 12 soil samples that could be mapped to an assembled protein sequence with a minimum sequence identity of 90% (step 1 above).

In **Supplementary Fig. 5b** we show the count of assembled amino acids within various coverage ranges. Coverage of an assembled protein sequence is the sum of the number of residues aligned to that sequence during mapping divided by the length of the assembled protein sequence.

Around 5 to 10 times more reads can be mapped to the set of protein sequences assembled by Plass (red) than to the set predicted by Prodigal on the Megahit assembly. The gains are particularly high for high coverages.

### Soil Reference Catalog assembly and analysis

For the Soil Reference Catalog (SRC), we downloaded from the sequence read archive (SRA) at the NCBI/NIH all 640 metagenomic datasets that (1) had the “soil metagenome” taxon identifier, (2) had dates between 01/2014 and 02/2018, (3) were sequenced on Illumina HiSeq or NovaSeq machines, and (4) had paired-end reads of at least 2 × 150 bp length. Sample identifiers are contained in a file `SRC_sample_ids.txt at` https://github.com/martin-steinegger/plass-analysis

Plass assembled the 18 Tbp of raw reads on a small cluster of servers with 2 × 8-core Intel Xeon E5-2640v3 CPUs and 128 GB RAM. We removed protein sequences shorter than 100 residues and redundancy-filtered the protein sequences from each sample using Linclust with options `--min-seq-id 0.95 --alignment-mode 3 −c 0.99 --cov-mode 1 --cluster-mode 2`). We pooled these 12 billion protein sequences and further reduced their redundancy by clustering with Linclust (`--cov -mode 1 -c 0.9 --min-seq-id 0.9`). The clustering was done hierarchically, since Linclust can only process 2^32^ — 1 sequences at once. The final set contains 2 022 891 389 sequences.

We chose two metagenomic soil sets (`SRR5919294` and `SRR6201924`) that were not part of the 640 datasets used for building the SRC. We merged overlapping read pairs using FLASH [23], sampled 100 000 merged reads per sample, predicted protein sequence fragments using Prodigal [34], and searched through the 90%-redundancy-filtered versions of SRC and the Uniprot database [28] using the `mmseqs map` workflow (see below). We computed the fraction of mapped reads out of the total read count while demanding a minimum sequence identity of 50% or 90% using the option `--min-seq-id`.

### Read mapping

In this study, we use the novel `mmseqs` map workflow from the MMseqs2 package to find very similar protein sequence matches in a protein sequence database. It first calls the `mmseqs prefilter` module (with a low sensitivity setting of `-s 2`) to detect high scoring diagonals and then computes an ungapped alignment using the `mmseqs rescorediagonal` module. In contrast to the mmseqs search workflow, for maximum speed no gapped alignment is computed, query sequences are not masked for low complexity regions (--mas*k*-mode 0), and no compositional bias correction is applied (`--comp-bias-corr 0`). By default, the mapping workflow requires that 90% of query sequence residues are aligned to a database sequence (`--cov-mode 2 −c 0.9`).

### Software versions used

We used the following version of software in this article, Prodigal `V2.6.3`, FLASH `v1.2.11`, Velvet `1.2.10`, SFA-SPA `0.2.1`, metaSPAdes `v3.10.1`, Megahit `v1.1.1-2-g02102e1`, eggnog-mapper `1.0.3`.

## Assembled protein sequence sets

The assembled protein sequence sets are available as FASTA formatted files at https://plass.mmseqs.org.

## Code availability

Plass is GPLv3-licensed open source software. The source code and binaries for Plass can be downloaded at https://github.com/soedinglab/plass.

## Data availability

All scripts and benchmark data including command-line parameters necessary to reproduce the benchmark and analysis results presented are available at https://github.com/martin-steinegger/plass-analysis.

## References

[1] Howe, A. C. et al. Proc. Natl. Acad. Sci. U.S.A. 111, 4904–4909 (2014).

[2] Li, J. et al. Nat. Biotechnol. 32, 834–841 (2014).

[3] Sunagawa, S. et al. Science 348 (2015).

[4] Xiao, L. et al. Nat. Biotechnol. 33, 1103–1108 (2015).

[5] Nielsen, H. B. et al. Nat. Biotechnol. 32, 822–828 (2014).

[6] Forslund, K. et al. Nature 528, 262 (2015).

[7] Zerbino, D. & Birney, E. Genome Res. 18, 821–829 (2008).

[8] Li, D. et al. Bioinformatics 31, 1674–1676 (2015).

[9] Nurk, S. et al. Genome Res. 27, 824–834 (2017).

[10] Ye, Y. & Tang, H. J. Bioinform. Comput. Biol. 7, 455–471 (2009).

[11] Yang, Y. et al. Bioinformatics 31, 1833–1835 (2015).

[12] Steinegger, M. & Söding, J. Nat. Commun. 9, 2542 (2018).

[13] Sczyrba, A. et al. Nat. Methods 14, 1063–1071 (2017).

[14] Kashtan, N. et al. Science 344, 416–420 (2014).

[15] Berube, P. M. et al. Scientific Data 5, 180154 (2018).

[16] Hyatt, D. et al. BMC Bioinformatics 11, 119 (2010).

[17] Vollmers, J. et al. PloS one 12, e0169662 (2017).

[18] van der Walt, A. J. et al. BMC Genomics 18, 521 (2017).

[19] Huerta-Cepas, J. et al. Mol Biol Evol 34, 2115–2122 (2017).

[20] Tian, W. & Skolnick, J. Journal of molecular biology 333, 863–882 (2003).

[21] Lee, S. T. M. et al. Microbiome 5, 50 (2017).

[22] Carradec, Q. et al. Nat. Commun. 9, 373 (2018).

[23] Hingamp, P. et al. ISME J. 7, 1678–1695 (2013).

[24] Steinegger, M. & Söding, J. Nat. Biotechnol. 35, 1026–1028 (2017).

[25] Ovchinnikov, S. et al. Science 355, 294–298 (2017).

## References

[23] Magoc, T. & Salzberg, S. L. Bioinformatics 27, 2957–2963 (2011).

[24] Ye, Y. & Tang, H. J. Bioinform. Comput. Biol. 7, 455–471 (2009).

[25] Yang, Y. et al. Bioinformatics 31, 1833–1835 (2015).

[26] Steinegger, M. & Söding, J. Nat. Commun. 9, 2542 (2018).

[27] Sheetlin, S. et al. Bioinformatics 32, 304–305 (2016).

[28] Mirdita, M. et al. Nucleic Acids Res. 45, D170–D176 (2017).

[29] Kanehisa, M. et al. Nucleic Acids Res. 45, D353–D361 (2016).

[30] Steinegger, M. & Söding, J. Nat. Biotechnol. 35, 1026–1028 (2017).

[31] Sczyrba, A. et al. Nat. Methods 14, 1063–1071 (2017).

[32] Kashtan, N. et al. Science 344, 416–420 (2014).

[33] Berube, P. M. et al. Scientific Data 5, 180154 (2018).

[34] Hyatt, D. et al. BMC Bioinformatics 11, 119 (2010).

[35] Huerta-Cepas, J. et al. Mol Biol Evol 34, 2115–2122 (2017).

[36] Huerta-Cepas, J. et al. Nucleic Acids Res 44, D286–93 (2016).

[37] Lee, S. T. M. et al. Microbiome 5, 50 (2017).

[38] Carradec, Q. et al. Nat. Commun. 9, 373 (2018).

[39] Frith, M. C. Nucleic Acids Res. 39, e23 (2011).

[40] Hingamp, P. et al. ISME J. 7, 1678–1695 (2013).

