## Supplementary material for "Protein-level assembly increases protein sequence recovery from metagenomic samples manyfold"

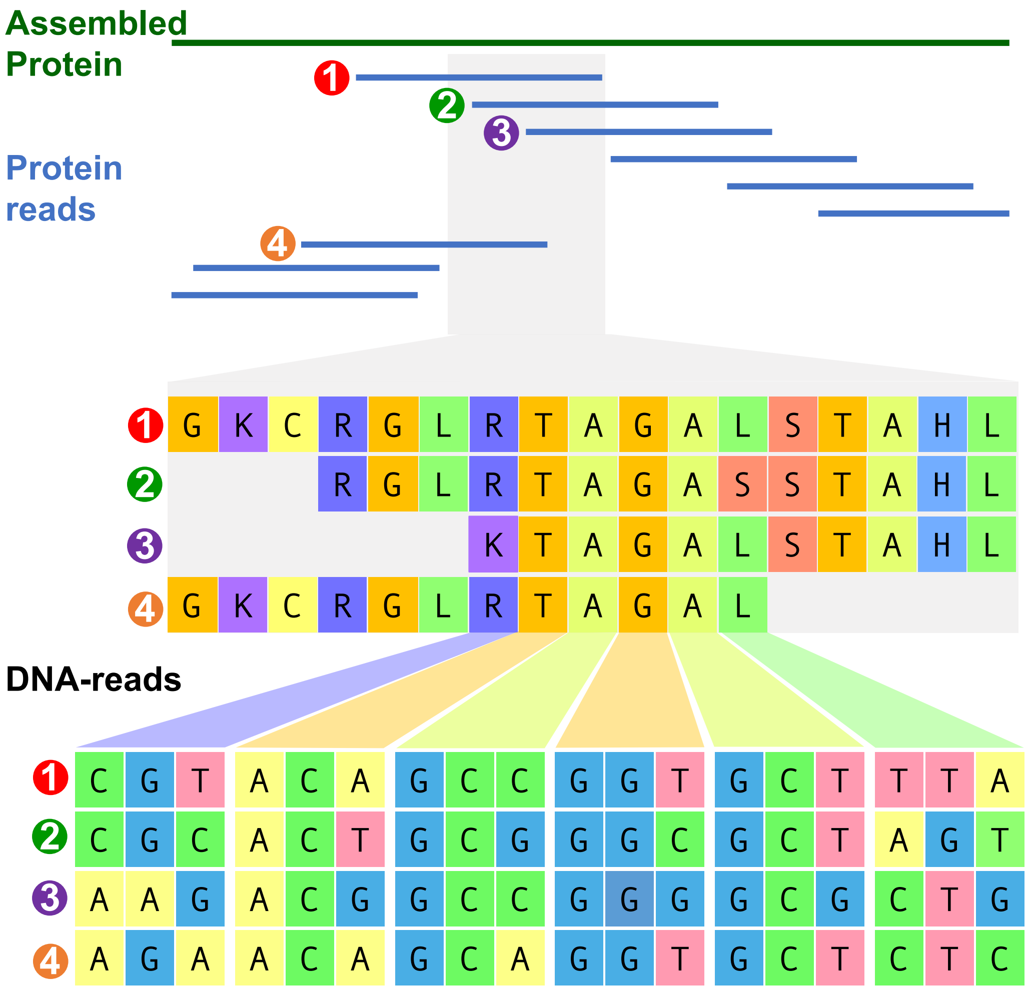


**Supplementary Figure** 1. **Schematic comparison of a nucleotide- and a protein assembly.** On top is the final protein assembly followed by the stacked overlapping protein reads. The small gray section highlights the multiple protein sequence alignment of the overlapping reads and below the respective nucleotide alignment. Less ambiguity is visible on the protein level due to conservative mutations (mutations with similar biochemical properties) compared to the nucleotide level, resulting in an assembly that is more robust to microdiversity in the population.

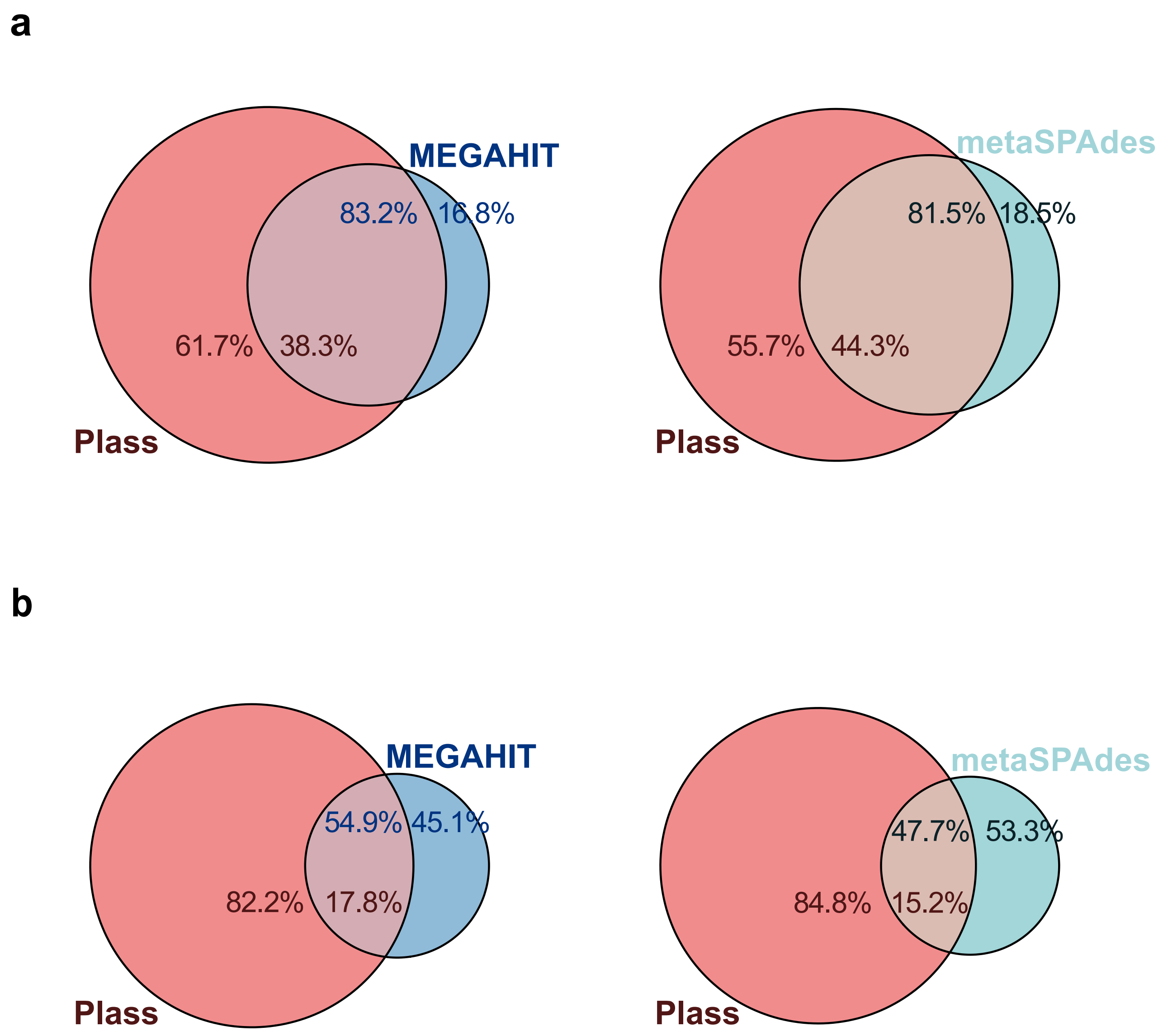


**Supplementary Figure** 2. **Overlap of assemblies of Plass with Megahit and metaSPAdes.** (**a**) left: A fraction 38.3% of amino acids in the Plass-assembled proteins of set 1 is covered by alignments to proteins in the Megahit assembly at a minimum sequence identity cut-off of 99%. Conversely, 83.2% of proteins in the Megahit-assembled set 1 is covered by alignments with Plass-assembled proteins. Right: Same as on the left but comparing the Plass assembly with the metaSPAdes assembly. (**b**) Same as (a) but for protein set 2.

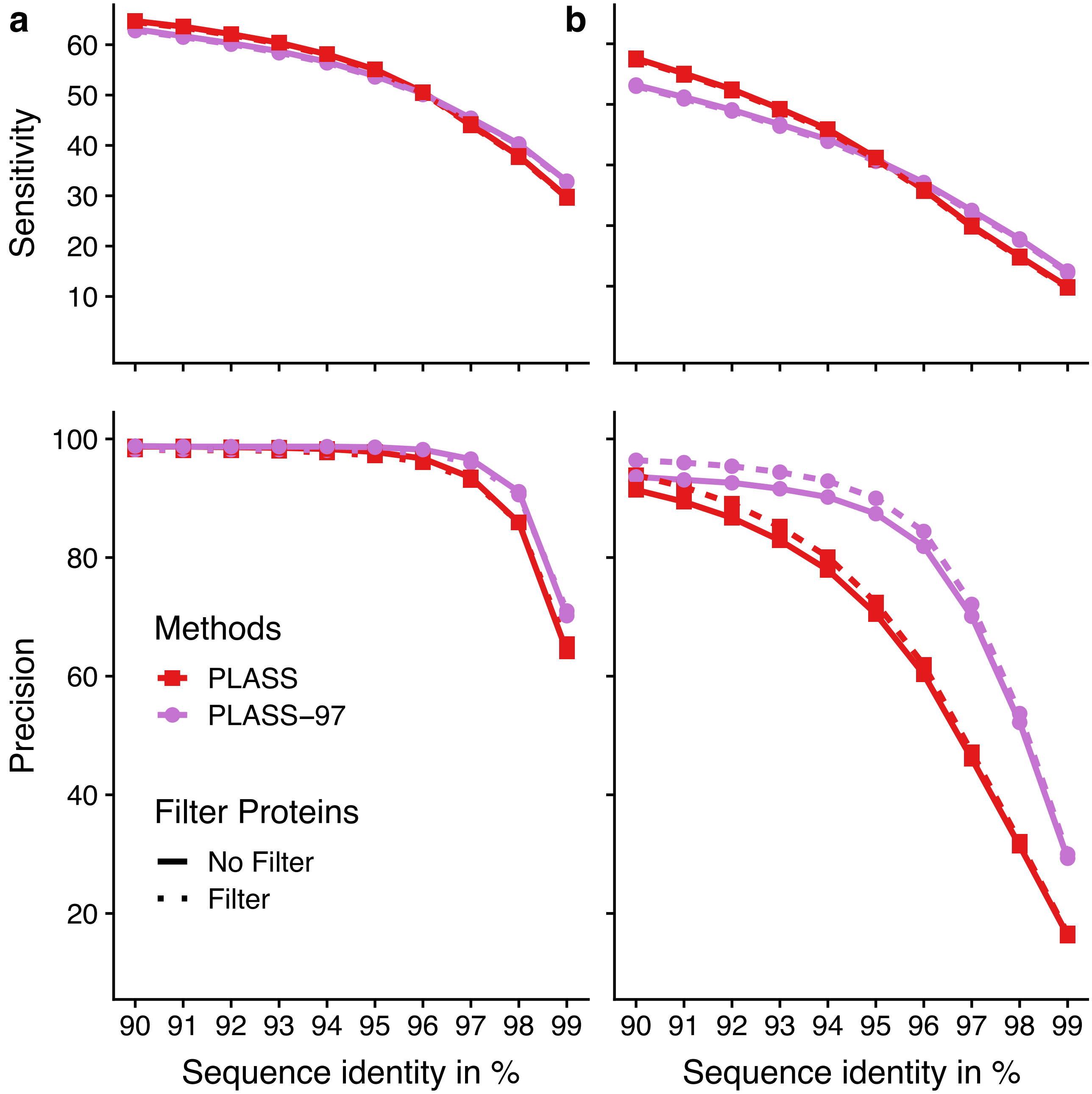


**Supplementary Figure 3. Effect of neural network filter to remove wrong translation frames.** Sensitivity and precision in set 1 (**a**) and set 2 (**b**). Top: assembly sensitivity is the fraction of reference sequence amino acids that matches to an assembled protein sequence. Bottom: assembly precision is the fraction of assembled amino acids that matches to a reference protein at the minimum sequence identity on the $x$-axis. Plass uses a minimum sequence identity for merging fragments of 90%, Plass-97 uses a threshold of 97%.

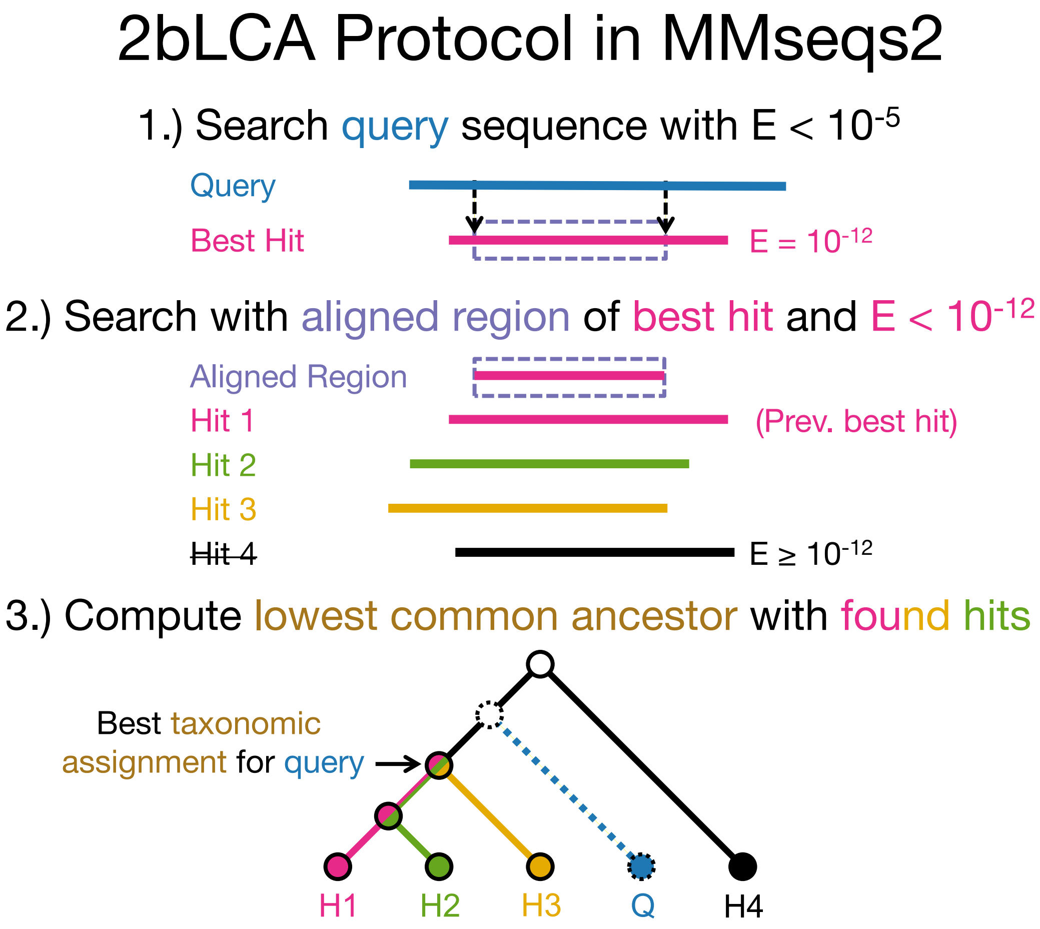


**Supplementary Figure** 4. **Comparison of Megahit assignment using the 2bLCA protocol.** The MMseqs2 taxonomy assignment workflow uses three steps to assign a taxonomic label to a query sequence. (1) We search with the query sequence against a reference database and extract the aligned subsequence of the best hit. (2) This sequence is matched again against the reference database. Each hit with an $E$-value smaller than the best hit $E$-value from the previous search is accepted. (3) We compute the lowest common ancestor based on the taxonomic labels of all accepted hits.

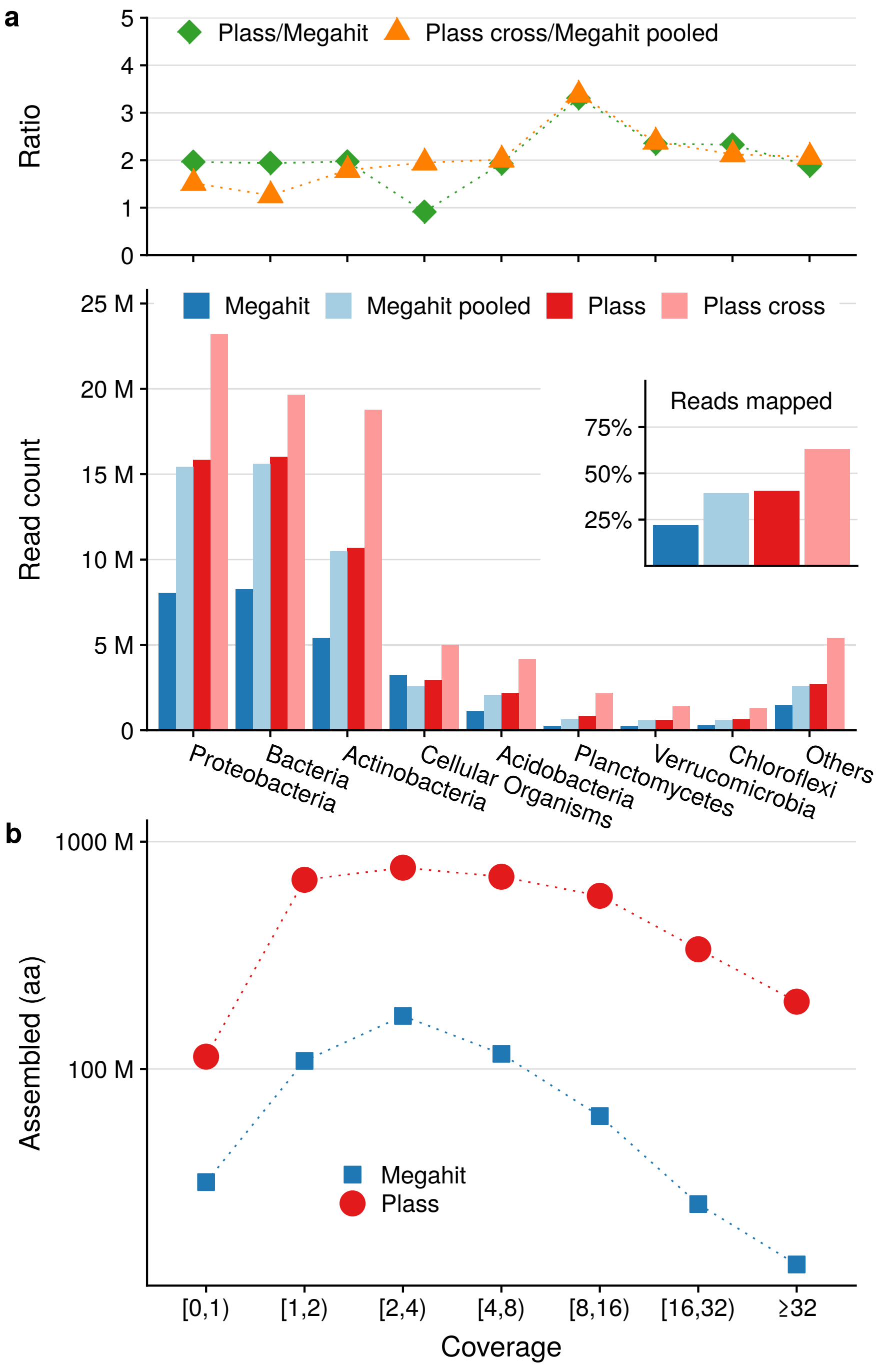


**Supplementary Figure** 5. **Taxonomy evaluation of the soil metagenome assembly.** (**a**) We investigate the taxonomic composition of the 8 most abundant taxa (all other taxa are pooled in “Others”) in the soil assemblies from **Fig. 2d** (blue: Megahit, red: Plass) and the assemblies of the 12 soil samples from **Fig. 2e** (light blue: Megahit, light red: Plass). On top we show the read count ratios between Plass and Megahit, for both the single and 12 soil assemblies. The inset gives the fraction of reads in the single and the 12 soil samples that could be mapped to an assembled protein sequence. (**b**) We show the count of assembled amino acids within various coverage ranges for Megahit (blue) and Plass (red) in the single soil sample.

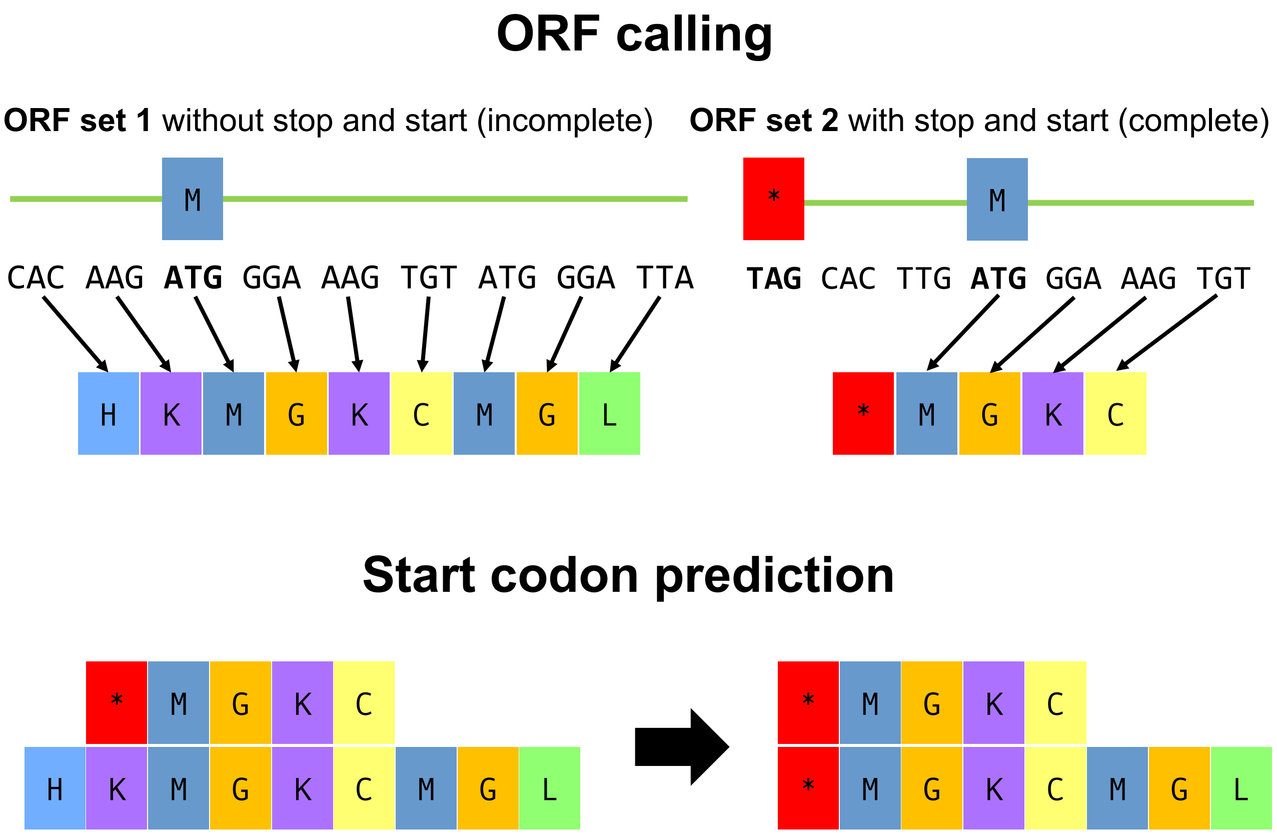


**Supplementary Figure** 6. **Plass ORF extraction and start codon prediction** )**ORF calling**) Plass extracts two sets of ORFs. ORF set 1 contains all translated ORFs with at least 45 codons. ORF set 2 contains all translated ORFs with at least 20 codons starting with a putative `ATG` start codon that is the first `ATG` codon after a stop codon in the same frame. (**Start codon prediction**) Plass predicts start codons with a consensus method using a multiple sequence alignment of ORF set 1 and 2. Wherever at least 20% of all methionines in one column are marked by a prepended asterisk, it removes the preceding residues from all other sequences and prepends an asterisk to all sequences to mark the start.
